# Convergent evolution of venom gland transcriptomes across Metazoa

**DOI:** 10.1101/2021.07.04.451048

**Authors:** Giulia Zancolli, Maarten Reijnders, Robert M. Waterhouse, Marc Robinson-Rechavi

**Author notes:** Corresponding author: Giulia Zancolli, **Email:**. **Author Contributions:** GZ and MRR conceived the study design. GZ collected all data and performed all analyses except where specified, with input from MRR and RMW; MR performed GO annotations and GO term semantic similarity analysis; RMW performed species phylogeny. GZ wrote the first draft of the manuscript, and GZ and MRR wrote the final version.

## Abstract

Animals have repeatedly evolved specialized organs and anatomical structures to produce and deliver a cocktail of potent bioactive molecules to subdue prey or predators – venom. This makes it one of the most widespread convergent functions in the animal kingdom. Whether animals have adopted the same genetic toolkit to evolved venom systems is a fascinating question that still eludes us. Here, we performed the first comparative analysis of venom gland transcriptomes from 20 venomous species spanning the main Metazoan lineages, to test whether different animals have independently adopted similar molecular mechanisms to perform the same function. We found a strong convergence in gene expression profiles, with venom glands being more similar to each other than to any other tissue from the same species, and their differences closely mirroring the species phylogeny. Although venom glands secrete some of the fastest evolving molecules (toxins), their gene expression does not evolve faster than evolutionarily older tissues. We found 15 venom gland specific gene modules enriched in endoplasmic reticulum stress and unfolded protein response pathways, indicating that animals have independently adopted stress response mechanisms to cope with mass production of toxins. This, in turn, activates regulatory networks for epithelial development, cell turnover and maintenance which seem composed of both convergent and lineage-specific factors, possibly reflecting the different developmental origins of venom glands. This study represents the first step towards an understanding of the molecular mechanisms underlying the repeated evolution of one of the most successful adaptive traits in the animal kingdom.

## Introduction

Organisms often evolve predictably similar features when presented separately with the same environmental or biological challenge (1). A long-standing question is whether the repeated evolution of adaptive traits in distinct lineages involves similar molecular changes, such as protein-coding sequences, cis-regulatory DNA elements, or gene expression (1–6). Animal venom represents one of the most remarkable examples of convergent evolution. On more than 100 occasions animal lineages have independently evolved the ability to secrete potent molecules to subdue prey or predators. Despite having the same biological role, the origin, anatomy and organization of the venom apparatus differ dramatically among lineages (7). Venom systems therefore represent an exceptional opportunity to test whether different animal lineages have repeatedly adopted similar molecular mechanisms to perform the same function (8).

Recent advances in sequencing technologies have allowed the molecular characterization of hundreds of proteomes and transcriptomes from the venom glands of several taxa, with a focus on medically important ones such as snakes and spiders, but also more neglected lineages (9–11). In particular, most venom gland transcriptome studies are focused on the identification of toxin transcript sequences, although recently there has been an increased interest for the “non-toxin” transcriptome (12, 13). The availability of genomes and of venom-gland RNA-seq datasets from various venomous lineages provide the opportunity for a comparative analysis across the animal tree of life to answer a simple, yet unexplored questions: did animals independently employ the same genetic toolkit to achieve the same function?

Here we address this question by comparing, for the first time, gene expression profiles from 20 venomous species representing eight independent origins of venom, and spanning ∼ 700 million years of evolution (protostome / deuterostome divergence) (14). We tested whether convergence in the ability to produce venom corresponds to the convergent evolution of gene expression levels in the venom glands. As venomous animals have evolved specialized organs for the biosynthesis and secretion of toxins, we expect enrichment of similar biological processes even among distantly-related taxa. However, animal venom glands are non-homologous structures with diverse origins (8); therefore, we hypothesize that convergence in biological function does not imply similarity in global gene expression profiles and regulatory networks. To study this, we first used a set of conserved ortholog genes to assess global evolutionary patterns of expression across all taxa. Then, we examined the whole transcriptome in each species separately to determine lineage-specific or shared expression changes, pathways and regulatory networks. We found a striking similarity of global gene expression patterns in evolutionary distinct venom glands, especially in genes involved in secretory functions, which indicates that complex trait evolution may sometimes be more constrained and predictable than expected. On the other hand, lineage specific profiles suggest that the way in which cells are regulated and communicates might reflect the diverse developmental origins of venom systems.

## Results and Discussion

### Does convergence in function correspond to convergence in gene expression profiles?

To analyze to what extent gene expression profiles of non-homologous venom glands are convergent, we compared publicly available RNA-seq datasets of venom glands and other body tissues from 20 venomous species representing eight different origins of venom: five spiders, two scorpions, one bee, three wasps, one fly, two mollusks, four snakes, one fish, and one mammal (Table 1 and *SI Appendix*, Table S1). In total, we used 2,528 orthogroups, sets of ortholog genes across all species, to create an expression matrix of log-transformed, quantile normalized Transcripts Per Million (TPM) values (*SI Appendix*, Dataset S1).

**Table 1.**
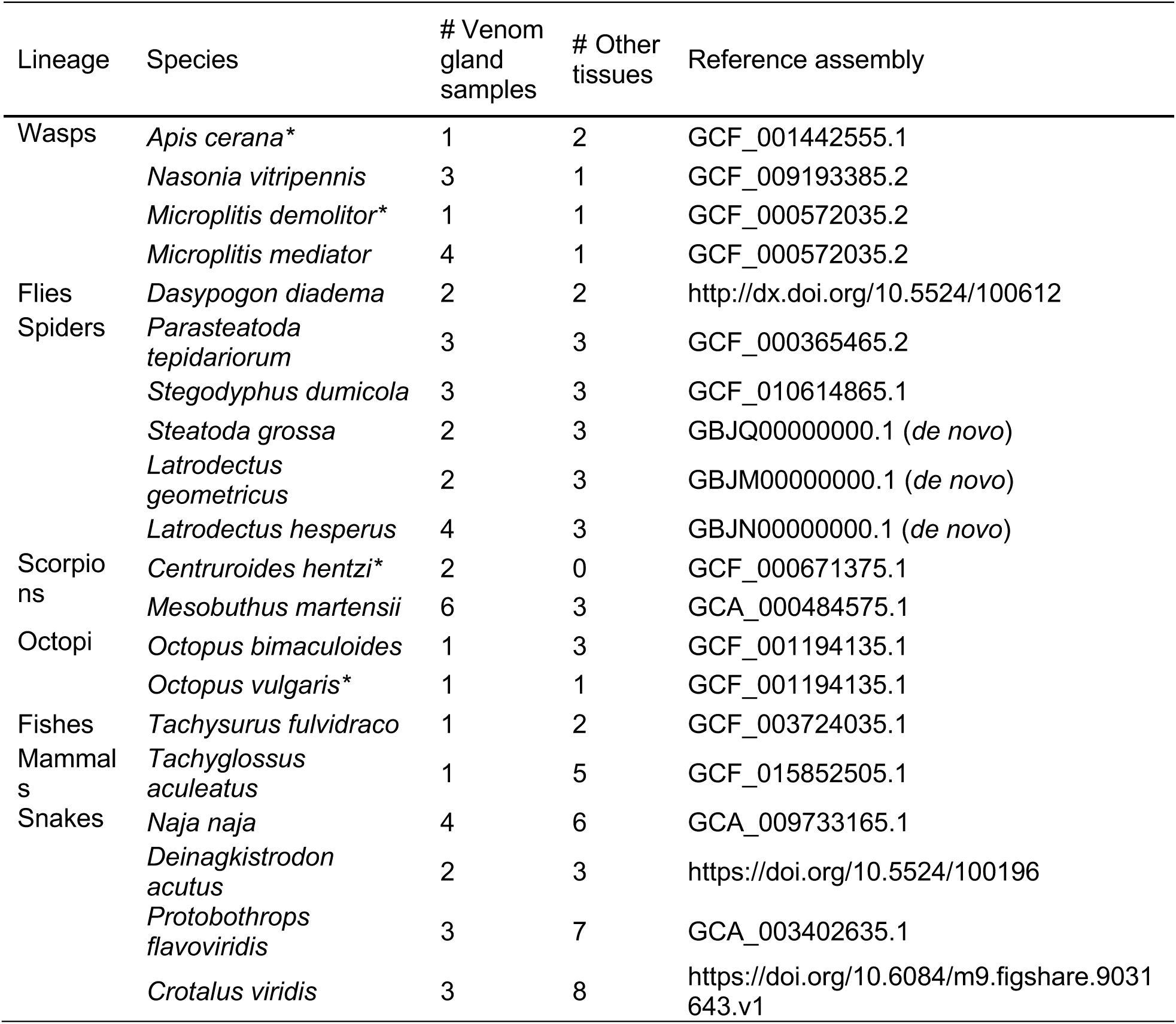
List of 20 species included in the analysis representing eight independent origins of venom, one per lineage. The table reports the number of venom gland samples, number of other tissues and the reference assemblies used for the analyses. Transcript and protein sequences were obtained either from the NCBI Genome (GCA and GCF) databases, the NCBI Transcriptome Shotgun Assembly (for *de novo* transcriptome assemblies), or from other repositories as indicated. *Species not used in the species-level differential expression analysis.

First, we explored gene expression patterns between lineages and tissues with principal component analysis (PCA), and found that the first three components clearly separated the venom gland samples from the other body tissues (Fig. 1 and *SI Appendix*, Fig S1). We had hypothesized that the expression levels of toxin genes, which are generally either restricted to or highly enriched in the venom gland, might be the drivers of this strong pattern. However, after removing the 38 orthogroups corresponding to known toxin genes from the expression matrix, the PCA did not substantially change (*SI Appendix*, Fig. S2). The clustering of venom gland samples in the PCA is remarkable considering the diverse origins of the datasets, not only because they are from different species and experiments, therefore different RNA processing and sequencing techniques, but also in terms of sampling procedures (e.g. time of dissection, pooling of samples, sex, age). Thus, it most likely represents a lower estimate of the degree of molecular convergence of the venom glands.

**Figure 1.**
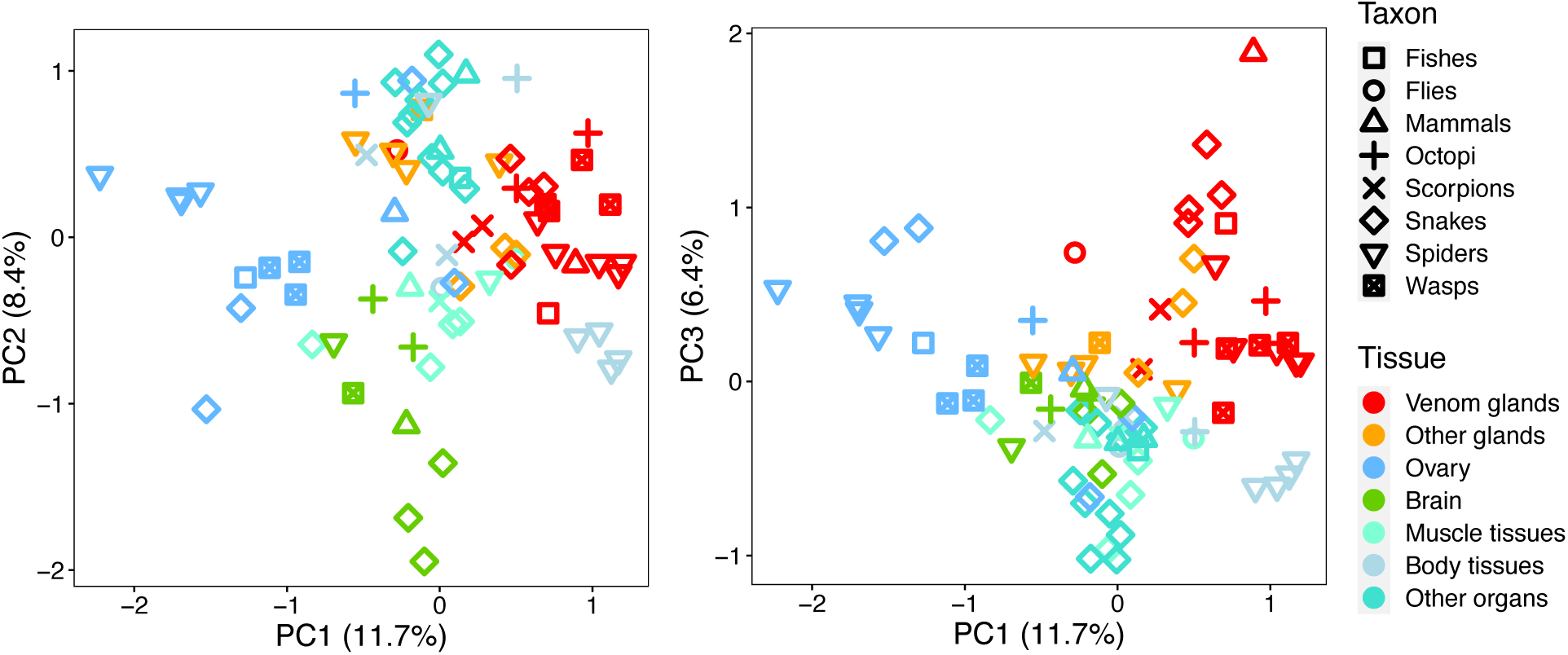
Global patterns of gene expression differences between multiple tissues and lineages. Principal component analysis based on the normalized expression levels (TPM) of 2,528 shared orthogroups among all taxa. The proportions of variance explained by the components are in parenthesis. ‘Body tissues’ include: abdomen, body tissue, cephalothorax, and viscera; ‘Other glands’ include: accessory venom gland, hypopharyngeal gland, rictal gland, salivary gland and silk gland; ‘Muscle tissues’ include: muscle, proboscis, heart, and leg; ‘Other organs’ include: liver, kidney, and pancreas.

Venom glands are mostly composed of epithelial secretory cells; therefore, we can expect their gene expression profiles to be similar to that of other exocrine tissues. Indeed, non-venom glandular tissues were positioned between the venom glands and the other tissues in the principal component space (Fig. 1 and *SI Appendix*, S1), suggesting shared expression patterns between secretory tissues. The rattlesnake’s accessory venom gland and the Indian cobra’s salivary gland clustered with venom glands: the accessory venom gland, as the name suggests, contributes to the production of venom and therefore it is expected to have a similar transcriptome (15). The pattern of clustering of the Indian Cobra’s salivary gland is more complex; one possible explanation is that the gland may have been mis-identified or there might have been contamination during dissection. The variation and diversity of secreting dental glands in snakes makes it difficult to clearly distinguish them, and confusion in the terminology of these homologous structures has been previously noted (16). Nonetheless, snake venom gland transcriptomes seem to rely on a conserved secretory gene regulatory network shared with salivary tissues of other amniotes (13).

Considering the diverse, and often recent, evolutionary independent origins of venom glands, we hypothesized that they would have higher transcriptome similarity to other tissues of the same species rather than between venom glands of different species. Contrary to this expectation, venom glands were more similar to each other than to any other tissue even from the same species (Spearman rank correlation coefficient ρ, Fig. 2). This is consistent with their clustering in the PCA (Fig. 1), although the comparisons were not statistically significant after Benjamini-Hochberg correction due to the low sample size of the non-venom tissues. However, we observed some particularly low correlation values between venom glands of the echidna and of other species. The echidna has a peculiar venom system compared to other animals – the venom gland is only active during the breeding season, it is found only in males, and it is thought to play a role in scent communication and to aid in competition. Furthermore, the loss of ability to erect the spur for venom delivery is thought to be the result of gradual decay of venom function (17). For these reasons, it might be that the echidna’s venom gland is diverging from the shared function of high-level secretory machinery in other animals, thus the observed low similarity values and the separation from the other tissues in the PC plot (Fig. 1).

**Figure 2.**
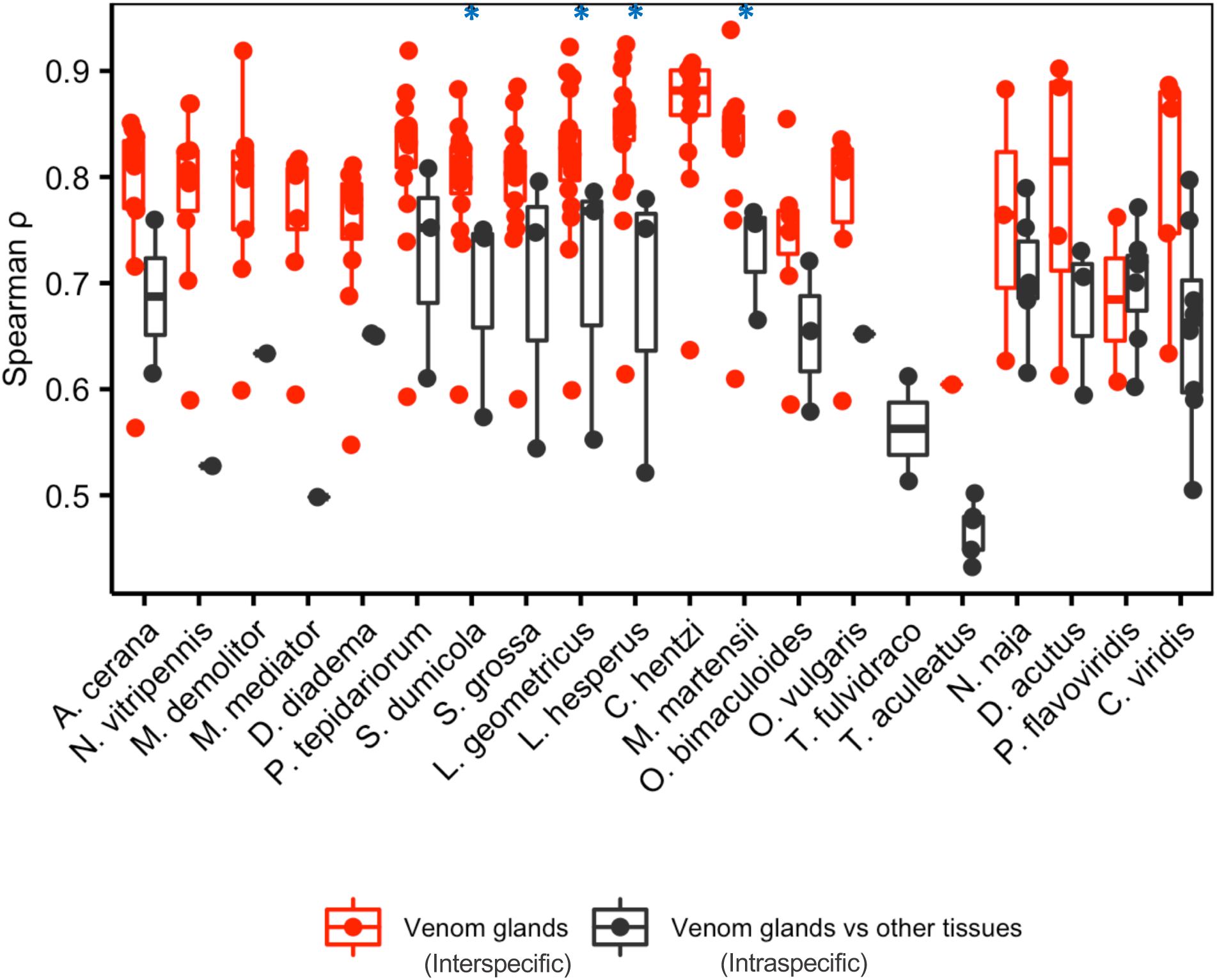
Transcriptome similarity between venom glands and other tissues. For each species, interspecific similarity (Spearman rank correlation coefficient ρ) between venom glands is compared to intraspecific coefficients between the venom gland and all other tissues for that species. Significant comparisons (Wilcoxon signed-rank test, *p* < 0.05 before correction) are indicated with an asterisk, although they were not significant after Benjamini-Hochberg correction. The low data points correspond to the pairwise correlations with echidna.

### Do closely related lineages have more similar transcriptomes?

We hypothesized that the expression levels of genes from homologous venom glands, i.e. from species that share a common venomous ancestor (e.g. snakes), follow a phylogenetic pattern, as regulatory changes accumulate over time. On the contrary, when comparing non-homologous transcriptomes, we might observe different, unpredictable patterns, with distantly related lineages clustering together due to functional convergence. To test this hypothesis, we compared a venom gland expression tree with the species phylogenetic tree. The expression tree (Neighbor-Joining) was constructed using two distance metrics, 1-Spearman coefficient and Euclidean distances, and the species tree was based on multisequence alignments of 1:1 orthogroups using RAxML (18). Surprisingly, the expression tree was overall consistent with the phylogenetic tree (Fig. 3A). The expression tree correctly separated protostomes and deuterostomes and the eight independent origins of venom. The branching patterns within these clades also broadly reflected the known phylogeny, except for the tree based on Euclidean distances which grouped Octopi with Ecdysozoa (*SI Appendix*, Fig. S3).

**Figure 3.**
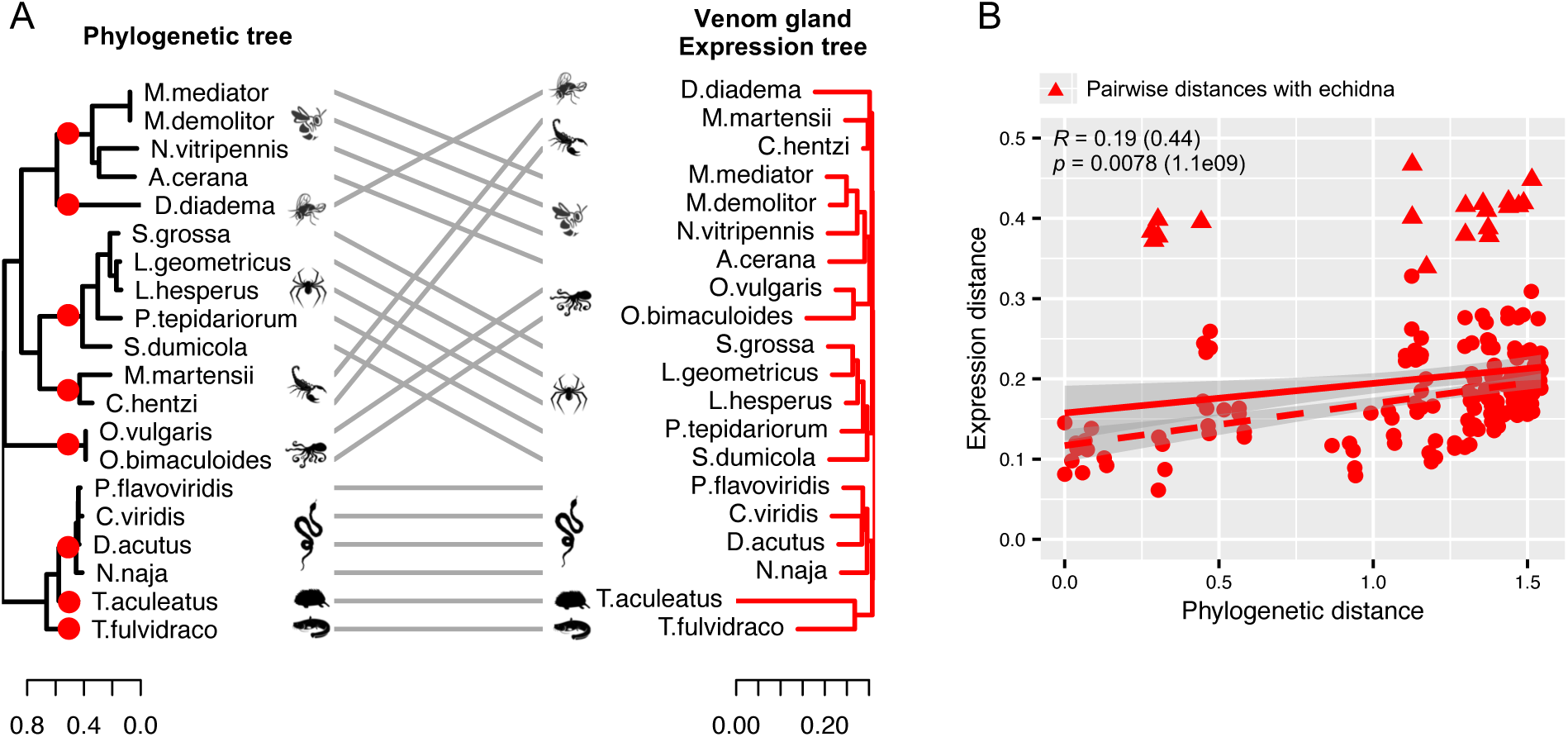
Comparison of venom transcriptomes and species phylogeny. A: Phylogenetic species tree (left), with circles marking the independent origins of venom in relation to venom gland expression tree (right) with 100% bootstrap support throughout (not shown). B: Sequence-based phylogenetic distances vs. venom gland expression distances (1-Spearman coefficient). Pair distances between echidna and the other species are marked with triangles, all the others are circles. The dotted line indicates the positive correlation between expression and phylogenetic distances excluding the echidna data points; the corresponding correlation test values are in parentheses.

Then, we compared sequence-based phylogenetic distances with expression distances between all species pairs to test whether closer lineages have more similar transcriptomes. Expression distances were positively correlated with phylogenetic distances (*R* = 0.19), confirming that closer taxa have more similar expression patterns (Fig. 3B). We noticed some particularly high expression distance values; these outliers are all pairwise distances between the echidna and other animals. As mentioned above, the echidna’s transcriptome is particularly divergent from all the other animals. The correlation between phylogenetic and expression distances was, as expected, much stronger excluding the echidna (*R* = 0.44, *SI Appendix*, Fig. S4).

### Do venom-gland transcriptomes evolve faster than those of other tissues?

Venom glands are derived traits that evolved from already differentiated tissues. Furthermore, the main product of venom glands, toxins, are among the fastest evolving genes, and their genomic makeup is highly variable and dynamic, with duplications and deletions between individuals of the same species (19). For these reasons we hypothesized that venom gland transcriptomes diverge at faster rates than tissues which are evolutionarily older. To test this hypothesis, we built gene expression trees based on 1-Spearman and Euclidean distances for other homologous tissues with at least seven species in our dataset, i.e. ovary and brain, and compared them with the venom gland expression tree (*SI Appendix*, Fig. S5).

Similarly to the venom gland tree, the brain tree separated deuterostomes and protostomes, and broadly reflected the species tree. On the other hand, the ovary tree resolved the two clades only with Euclidean distances, and even then, octopi clustered with the deuterostomes (*SI Appendix*, Fig. S6). Then, we compared pairwise phylogenetic distances with expression distances, and tested whether venom glands have higher evolutionary rates (i.e. higher slope values). Overall, at similar phylogenetic distances, ovary and brain transcriptomes were more divergent than venom gland transcriptomes (*SI Appendix*, Fig. S5); however, the venom gland slope was not steeper compared to that of the ovary (*F* = 0.41, *p* = 0.52), nor the brain (*F* = 1.41, *p* = 0.24). These results suggest that, contrary to our hypothesis, venom gland expression transcriptomes do not evolve faster than homologous, evolutionary older tissues, but they have comparable evolutionary rates.

### Are there venom-gland specific transcription modules?

The observed convergence between venom gland transcriptomes might be driven by a set of genes that have similar expression patterns, therefore producing tissue identity. We identified ‘modules’ by isa clustering (20) based on the expression matrix of all tissue types. In total, we found 62 modules (Fig. 4A and *SI Appendix*, Dataset S2 and S3), of which 36 included venom gland samples. Of these, 15 were exclusively venom-gland specific, 12 included venom glands and various glandular tissue samples, and the remaining nine were a mix of venom glands and other tissues. The venom gland-specific modules differed in species composition (5 to 17 species) and number of orthogroups (44 to 411 orthogroups); notably, modules 13, 24 and 40 included almost all species, thus representing a core gene set of 209 orthogroups. Some modules had lineage-enriched expression patterns, particularly for snakes; for instance, three modules included all snake species and one octopus, four modules included snakes and insect species, and one module included snakes and echidna. However, the genes in these snake-enriched modules were mostly also found in other venom-gland modules.

**Figure 4.**
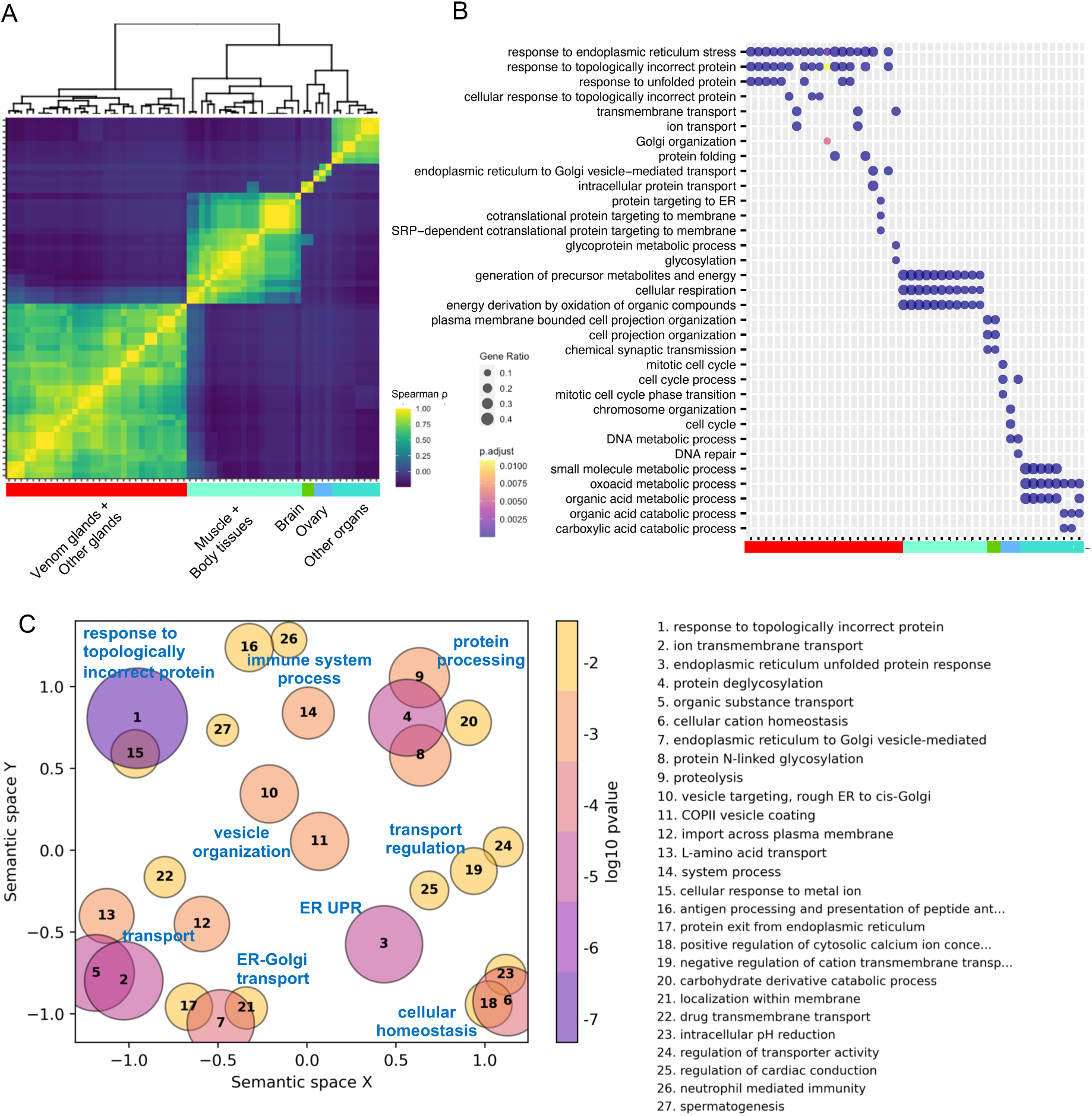
Tissue-specific modules and GO enrichment results. A: Heatmap based on Spearman correlation coefficients between the 62 modules; most modules are tissue-specific and cluster together. Definitions of tissue groups as in Fig. 1. B: Enrichment of the top 3 biological process GO terms of the tissue-specific modules. Color bar representing the tissues as in A. C: Visualization of biological process GO terms enriched in the venom gland core gene set (modules 13, 24 and 40) produced using GO-Figure! (59). Each bubble represents a cluster of similar GO terms summarized by a representative term reported in the legend, and sorted by the average p-values of the representative GO term across the three modules. Bubble size indicates the amount of GO terms in each cluster, and the color is the average p-value of the representative GO term across the gene modules. Similar clusters plot closer to each other.

Next, we screened for significant enrichment of KEGG pathways and GO functional categories based on annotation of human orthologs (Fig. 4B and *SI Appendix*, Dataset S4 and S5). Venom gland-specific modules were particularly enriched in pathways and in GO categories related to protein processing in endoplasmic reticulum (ER), secretion, transport, and particularly to stress response mechanisms (Fig. 4C). Similar results were obtained using *Drososophila melanogaster* orthologs (*SI Appendix*, Fig. S7). Also, we used TopAnat (21) to identify the anatomical structures where these gene modules are particularly expressed in an organism without venom glands. Genes belonging to venom gland specific modules were enriched in type B pancreatic cells and epithelial cells of pancreas in human, salivary glands and embryonic foregut in *D. melanogaster*. Modules which also included other glands were also enriched in entities of the oral and digestive system such as “mucosa of the sigmoid colon”, or “pylorus”. A link between venom glands and pancreas has been previously observed in snakes (8), and our results give a further hint that venom glands may have co-opted components from multiple anatomical origins. The orthogroups with the highest weight values in venom gland specific modules included genes involved in protein processing in ER and in ER stress, such as SEL1, SEC63, ERP44, DNAJC3, ER oxidoreductin 1, and SPCS3. Interestingly, the top orthogroups of the modules including venom glands and other glandular tissues were also related to protein secretion, such as UBA5, GRASP55/65, SRP54, and TM9SF, but they were not specifically related to ER or ER stress. The top orthogroups in the snake-enriched modules were mostly toxins or proteins also found in other venom gland modules. These findings emphasize the extreme secretory capacity of venom glands, and indicate that venom production necessitates the activation of stress response mechanisms. Dedicated protein-producing gland cells, such as those in venom-secreting tissue, have an exceptionally high secretory load relative to most cells. Consequently, during the emergence of venom gland cells, supporting mechanisms must have evolved to accommodate mass protein trafficking. One key mechanism is the unfolded protein response (UPR), which ensure reliable folding of proteins in the ER (22, 23). Our results suggest a central role of stress response mechanisms in enabling extreme cellular performance of venom glands, and this same mechanism seems to have been repeatedly adopted across the animal kingdom.

### Lineage-specific molecular mechanisms underlie convergence in general function of venom transcriptomes

The results so far point to convergence in gene expression profiles and the presence of venom gland-specific modules suggests concerted expression changes of genes involved in the secretory function. However, our first approach restricts the analysis to relatively few conserved orthogroups shared among all taxa, which might hinder the identification of lineage-specific patterns, or of patterns in fast-evolving genes missed by ortholog detection. To have a better resolution of the molecular underpinnings of venom gland activity, we performed differential expression and functional enrichment analyses for each species separately. For these species-level analyses, we retained only the 16 species for which we had multiple tissues and samples from the same study (Table 1), but we used all genes and enrichment analyses were based on each species’ annotation.

Across all species, the most upregulated genes, besides, of course, toxins, were involved in protein secretion and metabolism pathways (*SI Appendix*, Dataset S6). High level regulators of the unfolded protein response (UPR), e.g. ATF6, PERK, and IRE1, were upregulated in most lineages, confirming the isa-clustering results on orthogroups. Pathways that were enriched only in specific lineages were related to communication, such as “ECM-receptor interaction”, which was significantly upregulated in spiders, octopus and catfish, or “MAPK signaling pathway” and “GnRH signaling pathway” in all studied vertebrates (snakes, echidna, and catfish). A possible interpretation of these results is that the genes involved in the main function of the organ, i.e. protein secretion, are convergent, whereas the expression patterns of their regulators are inherited from developmental precursors, hence differences in signaling pathways might reflect the different origins of venom glands.

Enrichment analysis of GO biological process terms revealed that the most upregulated genes were mainly related to tissue development, regulation, signaling, transport, and metabolic and biosynthetic processes. A specific GO term was rarely found in more than one species, i.e. most GO terms were singletons; however, the enriched terms were semantically similar and grouped together (Fig. 5 and *SI Appendix*, Dataset S7). Additionally, some processes were found only in certain lineages, for instance reproduction and behavior were enriched only in scorpion, signaling pathways and nervous system in octopus, and the immune system in catfish. Membrane and ER were commonly enriched cellular components, while in some lineages extracellular region was also enriched.

**Figure 5.**
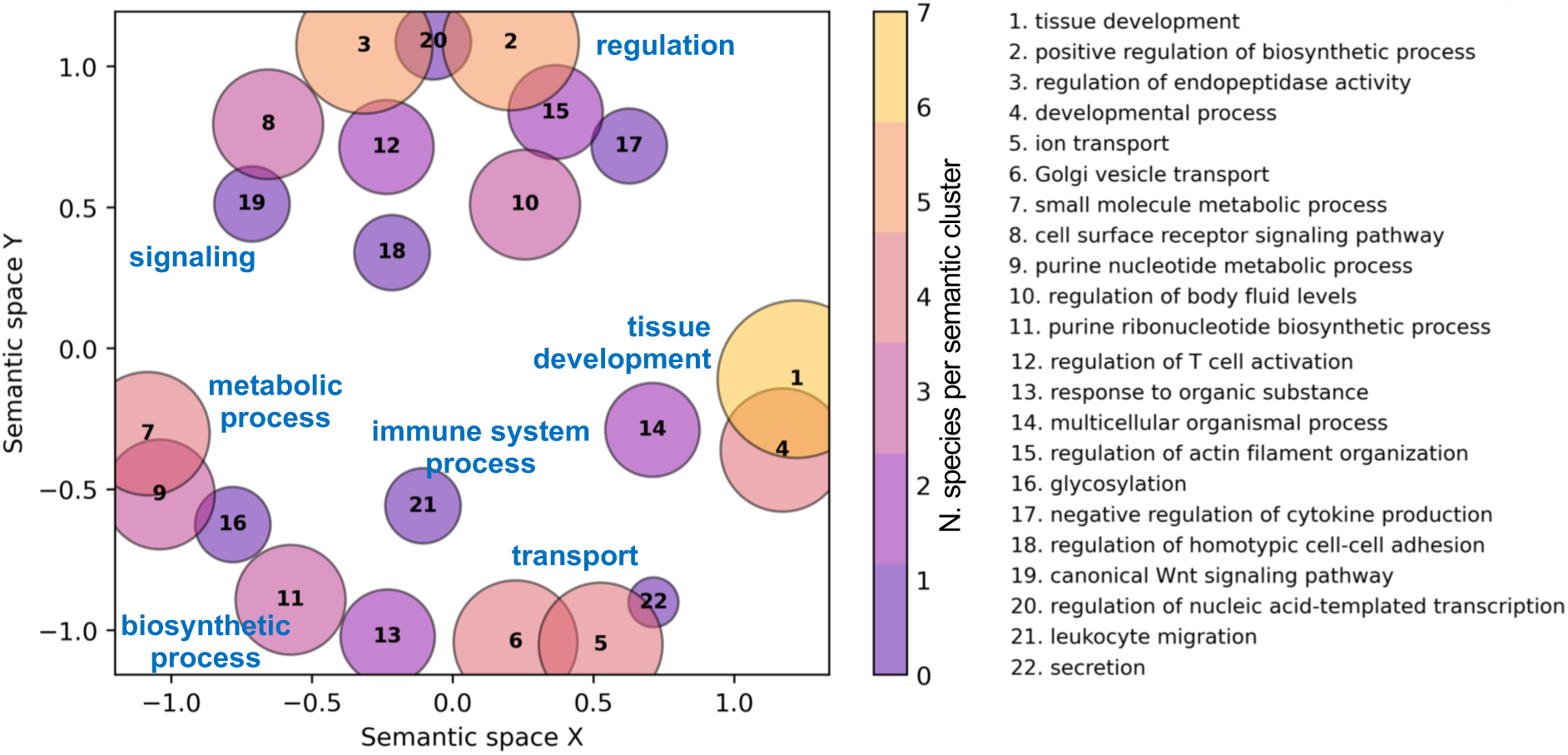
Semantic similarity scatterplot of GO biological process terms enriched in venom glands. Functional enrichment of upregulated genes was performed separately for each species, and all significant GO terms (p-value < 0.01) were summarized using GO-Figure! (59). Each bubble represents a cluster of similar GO terms summarized by a representative term reported in the legend and sorted by the amount of species with at least one term in the cluster. Bubble size indicates the number of terms in the cluster and the color corresponds to the number of species in the cluster. Similar clusters plot closer to each other.

The reasons why species were enriched with unique GO terms could be technical - GO annotations were done for each species separately, and this might have resulted in orthologous proteins being assigned to slightly different GO terms. Also, different sets of tissues were used in the differential expression analyses, and genome annotations vary in completeness between lineages. Nonetheless, our results suggest that there is an overlap in general biological processes, but that different lineages might have evolved different specific molecular mechanisms to perform the same general function. Even though global expression profiles are similar, the way in which biological processes are regulated might differ between species, hence convergence in function might not necessarily correspond to convergence in gene regulatory networks. To test this hypothesis, we examined which transcription factors (TFs) were upregulated (log_2_FC >1; FC = fold-change venom/average other tissues) in venom glands compared to other tissues. TFs were identified using the KEGG database and all the information related to the TFs were retrieved from the Human Protein Atlas (24), UniProt (25), and Bgee databases (21). The most upregulated TFs within each taxon were involved in ER stress and UPR response, in agreement with our previous results, but they were also largely involved in pluripotency, cell differentiation, and tissue development (*SI Appendix*, Dataset S8). We found thirteen orthologous TFs shared across all lineages, some of which were also found in venom gland-specific modules. These included TFs involved in the UPR and response to ER stress pathways (XBP1, CREB3), or typically expressed in the epithelium (ETS, BHLHB8/MIST1, BNC), but the majority were involved in cell proliferation, differentiation and growth (TWIST, MXD-MAD, FOXP2_4, SOX9). Various homeobox gene families that play pivotal roles in tissue development and differentiation were among the most upregulated TFs, but different members were found in different lineages: the NKX-homeodomain factor family, with NKX2.5 expressed in octopus, NKX1 in scorpions, and NKX3-1 in echidna and catfish; the SIX family, with SIX1 found in snakes and spiders while SIX4 in octopus, PBX, with PBX1 highly expressed exclusively in spiders and scorpions (Arachnida) and PBX3 in spiders and snakes, as well as DLX, and LHX. Myogenic factors were also upregulated, with MYOF5 found exclusively in spiders and MYOD1 in snakes (*SI Appendix*, Dataset S8).

An interesting finding was the high upregulation of the abdominal-B homeobox ABDB exclusively in scorpions and wasps. This TF specifies the identity of the posterior abdominal segments, the external genitalia and gonads, and is involved in regulating post-mating response. Another TF found only in these two lineages was the krueppel (KR) factor which is involved in differentiation of the Malpighian tubules, a type of excretory system in the posterior region of the alimentary canal of some arthropods. Compared to the other animals investigated, scorpions and wasps have in common the location of their venom apparatus which is in the posterior part of their body, or metasoma. In scorpions, venom glands are located in the telson where also cuticular pits and dermal gland openings have been described (26). The function of these other glands is not really understood, but it is hypothesized that they might produce sex pheromones and play a role in courtship (26). In parasitoid wasps, like those included in our study, venom glands are in the posterior dorsal surface and are connected to the female reproductive system. The close interaction between these two systems complements their functioning, since venom is injected through a modified ovipositor to ensure successful development of the offspring in the host. Several genes annotated with GO terms related to reproduction were expressed in the venom glands of both wasps and scorpions, and in the latter behavior and reproduction terms were significantly enriched (*SI Appendix*, Dataset S7). The exact function of these genes within venom glands is unknown. Nonetheless, these findings provide a first evidence that venom gland regulatory networks have evolved, to some extent, from the co-option of pre-existing genetic regulatory circuits of the tissues from which venom glands derive, or that are most closely related to.

The high number of genes and magnitude of enrichment related to cell cycle regulation in our results is intriguing, and might be indicative of high epithelial cell turnover. In many lineages, e.g. in scorpions (27), spiders (28), echidna (17), and catfish (29), venom is released by holocrine or apocrine modes of secretion, which cause cellular damage or complete destruction of the cell. Moreover, the high levels of cellular stress caused by massive toxin production might result in DNA damage, apoptosis, or cellular dysfunction. As a consequence, the activation of a regulatory network for epithelial cell turnover and maintenance might be necessary. Undifferentiated epithelial cells have been morphologically identified in the venom gland epithelium of various organisms, e.g. in spiders (30), scorpions (27), snakes (31). Furthermore, non-venom epithelial supporting cells and stromal cells, which express stem cell markers and niche factors, have been observed in snake venom gland single cell sequencing (31), supporting our finding of active cell growth and differentiation pathways in venom glands. These findings combined with our results suggest that conserved as well as lineage-specific regulators involved in cell differentiation and organ development have been repeatedly adopted during the evolution of venomous animals and that, besides secretion, regulation of cell cycle is a central task of venom glands.

## Conclusions

Many animal cell types possess the capacity for protein secretion, and a conserved molecular and organellar pathway exists for routing translated proteins out of the cell (23). However, dedicated protein-secreting cells, such as those producing venom, have an exceptionally high secretory load. During the evolutionary assembly of venom glands, stress response mechanisms seem to have been repeatedly adopted by different animals to cope with mass protein production. The resulting DNA damage, apoptosis and even complete destruction of those venom-producing cells with holocrine secretion, activate a regulatory network for epithelial development, cell turnover and maintenance. While sets of genes directly involved in the secretory function have similar expression patterns across animal lineages, and might thus be conserved, the way in which cells are regulated and communicates, are different between lineages, and might reflect their diverse developmental origins. Our findings provide a first evidence that venom gland regulatory networks have evolved, to some extent, from the co-option of pre-existing genetic regulatory circuits from the tissue most closely related to each venom glands. This study represents the first step towards an understanding of the molecular mechanisms underlying the convergent evolution of one of the most successful adaptive traits in the animal kingdom.

## Material and methods

### Species selection

For the analysis, we selected only venomous species with either an annotated genome or high quality *de novo* transcriptome available for the same species or a close relative, and with RNA-seq data of venom glands and other body tissues (Table 1 and *SI Appendix*, Table S1).

Considering the mixed nature of our dataset, for each species, we reduced the proteomes (proteins from annotated genomes and *de novo* transcriptomes) to a set of non-redundant sequences as follows: first, we filtered within-species 100% identical amino acid sequences with cdhit 4.6 (32) to select one representative sequence. Then, we compared with BlastP (33) the sequences against the NCBI non-redundant database (downloaded on 09.04.2020), the Uniprot-Toxprot (34) or Arachnoserver (35) databases, and retained only those with evalue < 1e-05. These processed proteomes were used for subsequent analyses.

### Orthogroup assignment

For each species, we assigned protein sequences to orthogroups using the mapping tool in OrthoDB v.10.1 (36). For the mapping we selected up to five closest taxa to assign proteins to orthogroups at the Metazoa node. In parallel, we compared the proteomes with BlastP against the same species selected for OrthoDB mapping. For each species, all Blast outputs were combined, and we kept only one hit per sequence (the one with the lowest evalue). Blast and OrthoDB mapping were then merged and proteins assigned to orthogroups using the OG2genes file at the Metazoa node. Species used for the orthogroup assignment are listed in *SI Appendix*, Table S2.

Orthogroups containing proteins reported as venom components in the reference genome/transcriptome paper, or that had the best Blast hit against a sequence in the Uniprot-Toxprot or Arachnoserver databases, were assigned as toxin-containing orthogroups.

### Expression levels

RNA-seq data were obtained from the Sequence Read Archive (SRA) database. Only Illumina SRA reads were selected; where possible, we selected at least three libraries for each tissue from each taxon, and only data generated from healthy, adult tissues were used. Raw fastq files were filtered with trimmomatic (37), their quality checked with fastQC (38), and quantified with kallisto (39) using default parameters for paired-end reads, and parameters -l 55 -s 1e-08 for single-end reads. All species were mapped to their own specific transcriptome with the exception of *Microplitis mediator*, which was mapped to *M. demolitor*, and *Octopus vulgaris* which was mapped to *O. bimaculoides*. We used *tximport* (40) to estimate transcript abundances as Transcript Per Million (TPM) for the metazoan-level analysis, and to aggregate read counts at the gene-level for the species-level analysis (see below).

### Orthogroup expression matrix

For the comparative analysis at the Metazoa node, orthogroup-level abundances were obtained as follows. Since most orthogroups included more than one protein per species (i.e. one-to-many or many-to-many orthologs), we selected one representative sequence for each orthogroup in each species as the transcript with the highest expression in venom gland samples (*SI Appendix* Dataset S9), and used the TPM value estimated for that transcript as the orthogroup expression value. All samples of all species were then merged into a matrix of orthogroup TPM values. To validate our method, we randomly selected one transcript per orthogroup, and we obtained similar results (see *SI Appendix*, Fig. S1 and S8).

Because the samples are from different experiments, to allow for comparison across samples, first, we minimized the effects of technical artifacts by quantile normalization on log_2_ transformed TPM values to which a pseudo count of 1 was added to prevent log_2_(0) scores. Then, we removed the batch effect caused by using multiple species and multiple SRA studies, using an empirical Bayes method implemented via the ComBat function in the sva R package (41) which has proven to be efficient with these kinds of datasets (13). Finally, we calculated tissue-level expression as mean TPM.

### Transcriptome similarity analysis

All analyses were performed in R version 3.6.2 (42). To obtain an overview of expression patterns we performed principal component analysis (PCA) with the rda function in *vegan* (43). To understand whether the observed pattern was biased by the shared expression levels of toxins in venom glands, we re-ran the analysis excluding the orthogroups containing toxin sequences. We quantified transcriptome similarity between tissues as Spearman rank correlation coefficients to test whether venom glands were more similar to each other than to any other tissue of the same species. Mean pairwise distances were compared with the Wilcoxon signed-rank test and p-values adjusted with (44).

### Phylogenetic tree

We constructed a phylogenetic species tree using 77 one-to-one orthologs. First, proteome redundancy was reduced by selecting a representative protein for all sequences > 90% identical using cd-hit 4.6. Then, we selected orthogroups that were single-copy in all species, or all except one. Orthogroup sequences were aligned with mafft 7.310 (45) and trimmed with trimAl 1.4.1 (46). The trimmed alignments were concatenated, trimmed and used to build the phylogenetic tree with raxml 8.2.12 (18), and bootstrap values of the consensus tree were obtained based on 100 replicates. The tree was rooted at the deuterostome-protostome split using the function root_in_edge of the R package *castor* (47).

### Expression trees

Expression trees were constructed for venom gland, ovary, and brain tissues. These homologous organs were chosen for the comparisons because they had the highest number of samples (minimum seven species).

We used the R package *ape* (48) to construct Neighbor-Joining expression trees based on two distance measures: 1 – Spearman rank correlation coefficient and Euclidean distances. The reliability of branching patterns was assessed with bootstrap analysis using 1000 replicates. When possible, trees were rooted at the deuterostome-protostome split. To verify whether closer taxa have more similar transcriptomes, we obtained pairwise distances using the function get_all_pairwise_distances in the package *castor*, and tested for correlation between phylogenetic and expression matrices with Mantel tests in the R package *vegan*. Expression divergence rates were calculated as the slope of linear regressions (lm) between pairwise expression distances and phylogenetic distances.

### Transcription Modules

We identified orthogroups with similar expression patterns using the iterative signature algorithm (isa) implemented in the *isa2* Bioconductor package (20) with default parameters. Briefly, isa identifies, in an unsupervised manner, sets of genes that exhibit coherent expression patterns over subsets of samples from large sets of expression data. It selects genes that are significantly under- or over-expressed in a random seed of samples, and then all samples are scored by the weighted average expression levels across these genes.

KEGG pathway and GO enrichment analyses of all isa modules were performed with *clusterProfiler* (49) based on human gene annotation after converting the OrthoDB ClusterId to NCBI EntrezId using the OG2gene file at the Metazoa node obtained from the OrthoDB data page. For both analyses, the foreground genes were the orthogroups of a module and the background genes were all the 2528 shared orthogroups.

For enrichment of anatomical structures, we used the TopAnat tool in Bgee 14.2 (21). TopAnat is based on TopGO (50) and works similar to a GO enrichment test except that it analyzes Uberon anatomical identities (51) where genes are expressed instead of GO terms. To find the anatomical entities in which tissue-specific gene modules were enriched, we run TopAnat based on human annotation and with the weight algorithm.

### Species-level differential expression analysis

To identify taxon-specific patterns, we performed differential expression analysis at the gene-level for each species separately using *edgeR* (52). Genes with low expression were automatically excluded with the function filterByExpr. Differences in library size were accounted for using TMM-normalization factors. We fit a quasi-likelihood (QL) negative binomial generalized log-linear model (glmQLFit function) to estimate empirical Bayes moderated dispersion; this method accounts for gene-specific variability from both biological and technical sources (53). Empirical Bayes QL F-tests were used to compare gene expression levels between venom glands and the average of the other tissue types. Genes with false discovery rate < 0.05 and log_2_ fold change > 1 were considered significantly upregulated.

KEGG pathway enrichment analysis of upregulated genes was performed with *clusterProfiler*. For the species which were not in the list of supported organisms in the KEGG catalog, we used the annotation of the closest species. Orthologs between the two species were identified by reciprocal BlastP.

As none of the species in this study have available GO annotations, we annotated all the proteomes using the program CrowdGO (54). CrowdGO is a consensus-based GO term meta-predictor that employs machine learning models combined with GO term semantic similarities and information contents to leverage strengths of individual predictors and produce comprehensive and accurate gene functional annotations. For the GO annotations in this study, we used the CrowdGOFull model which utilizes annotations from DeepGOPlus (55), FunFams (56), InterProScan (57), and Wei2GO (58).

GO enrichment analyses for the biological processes, molecular functions and cellular compartments were performed with TopGO (50) using the elim algorithm and fisher statistic test. We summarized the most enriched processes (pvalue < 0.01) across all species with GO-Figure! which reduces redundancy by grouping together GO terms with similar functions, and produces semantic similarity scatterplots where representative terms are plotted (59).

### Transcription factors analysis

Finally, we focused on expression patterns of transcription factors (TFs) to test for convergence in gene regulatory networks. We examined which transcription factors were upregulated in venom glands compared to other tissues, and whether these TFs were expressed across all species or only within certain lineages. For each species we downloaded the annotations from the KEGG BRITE database (‘03000 Transcription factors’). For species which were not included in the KEGG Organism catalog, we used the annotation of the closest species (as we did for the KEGG enrichment analysis). Because one KEGG orthology entry can contain multiple genes (similarly to the OrthoDB orthogroups), for the comparative analysis we kept one representative gene per KEGG ortholog.

## Acknowledgments and funding sources

This project has received funding from the European Union’s Horizon 2020 research and innovation programme under the Marie Skłodowska-Curie grant agreement No 845674 to GZ, and the Swiss National Science Foundation grant to PP00P3_170664 (RMW). We thank Nadia Ayoub for providing the amino acid sequences of the spider de novo transcriptomes, Robert Haney Zong-ji Wang for providing toxin sequences, and MRR lab members for bioinformatics support.

## Notes

### Competing Interest Statement

The authors have declared no competing interest.

### Summary of Updates

Figure 1 revised - added plot with PC1/PC2 Figure 4 moved to SI

