## Supplementary Information for "Convergent evolution of venom gland transcriptomes across Metazoa"

*Corresponding author: Giulia Zancolli

**This PDF file includes:**

Figures S1 to S8
Tables S1 to S2
Legends for Datasets S1 to S9

**Other supplementary materials for this manuscript include the following:**

Datasets S1 to S9

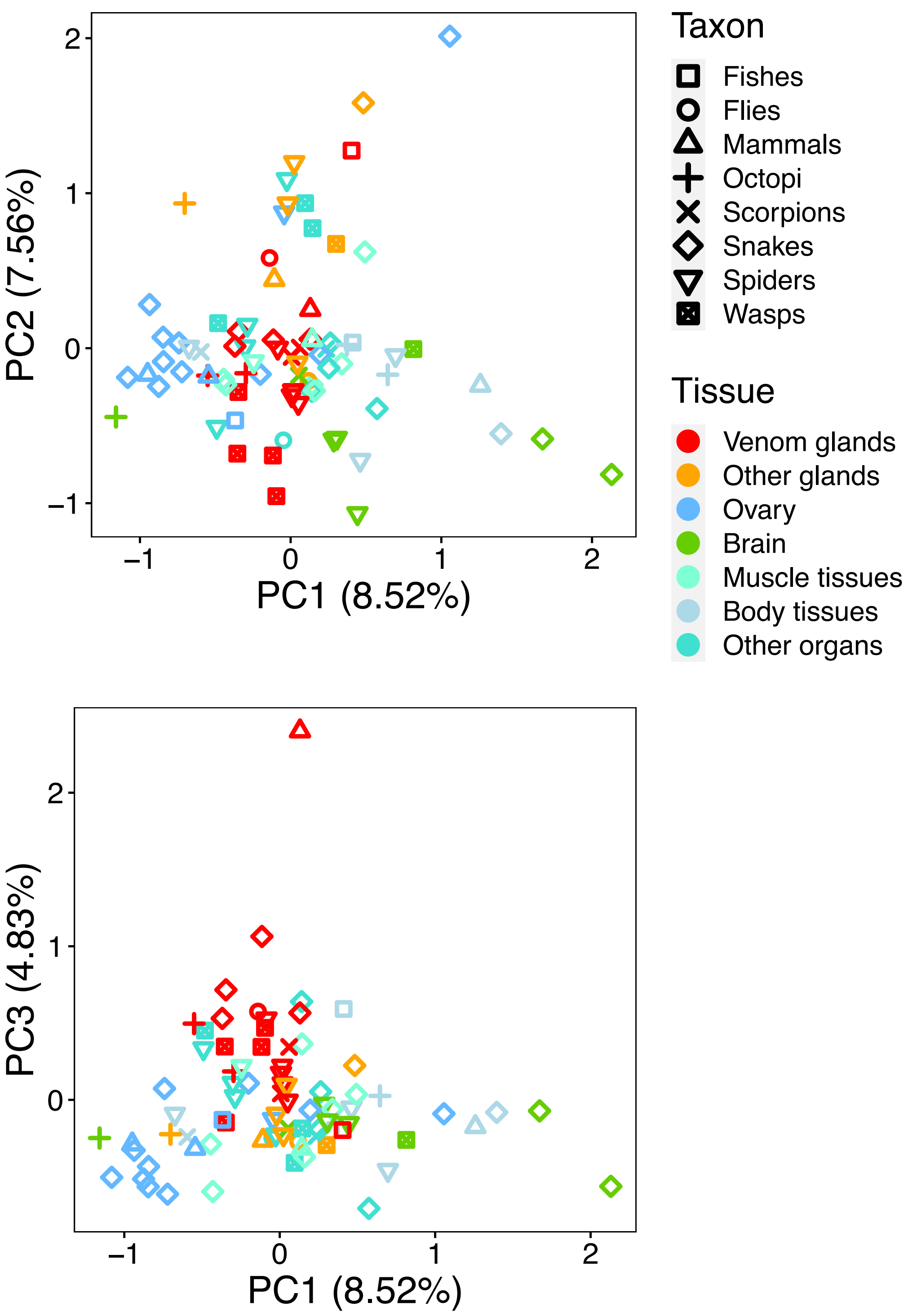


**Figure S1.** PCA using the expression matrix based on transcripts selected randomly for each orthogroup.

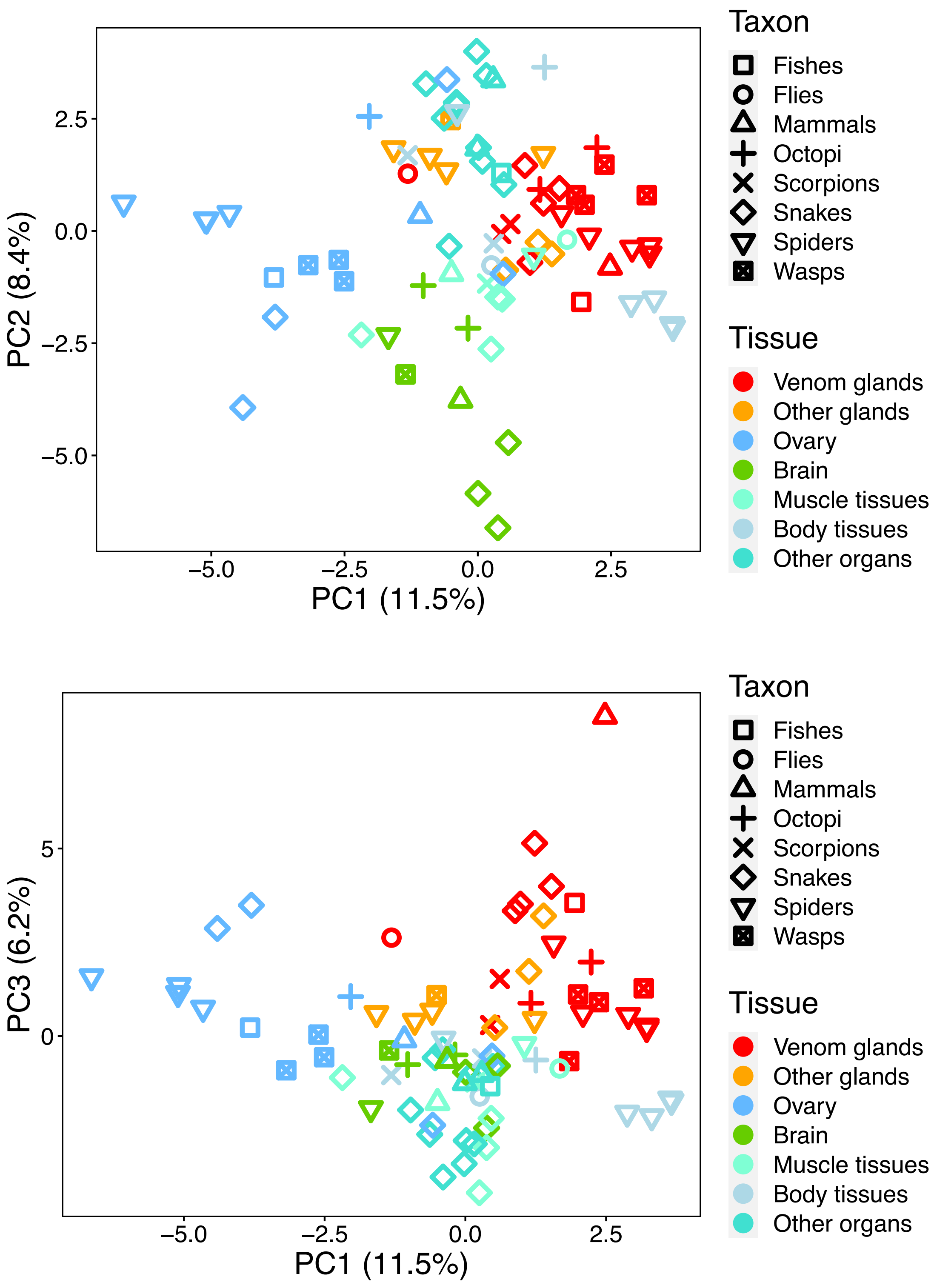


**Figure S2.** Principal component analysis based on 2,490 shared orthogroups. Orthogroups containing toxin sequences were excluded from the expression matrix.

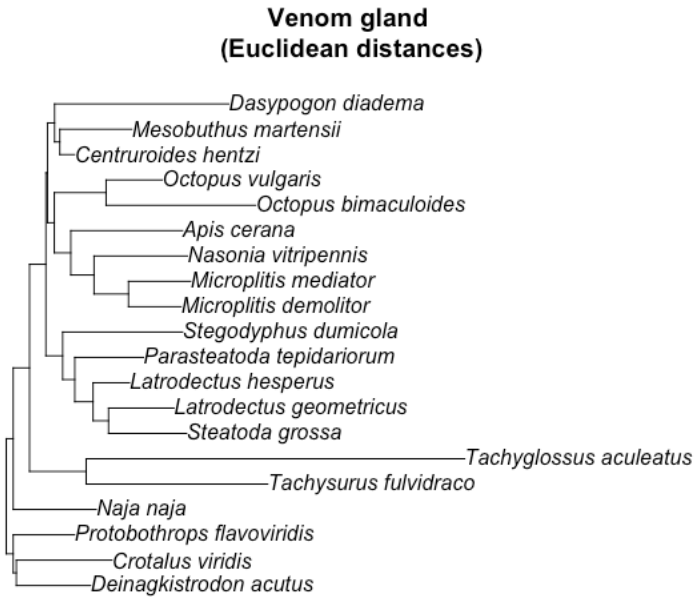


**Figure S3.** Venom gland expression tree based on Euclidean distance matrix. Overall the expression tree reflects the phylogenetic tree (Fig. 3 main text), except for the octopi which cluster with wasps.

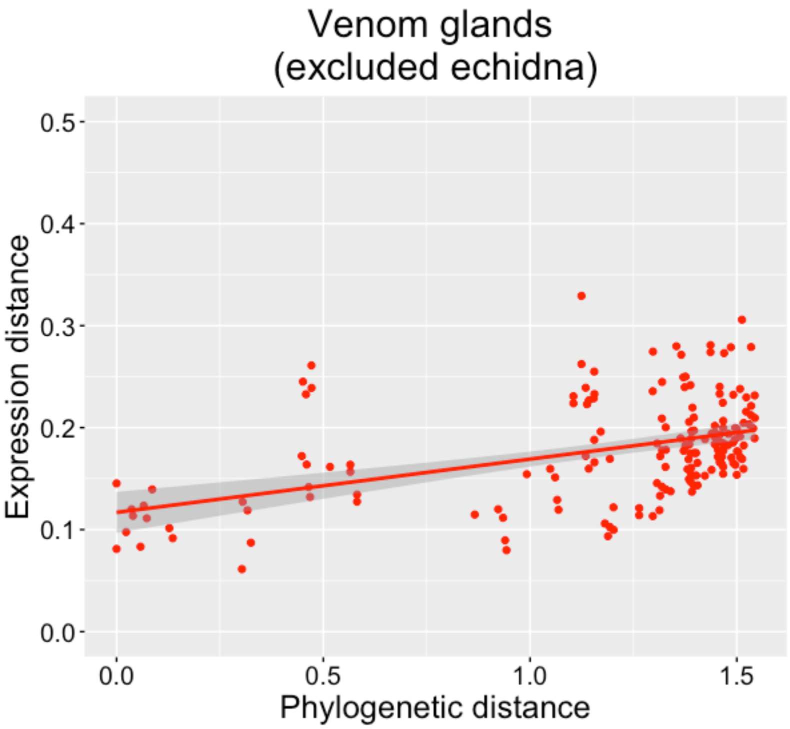


**Figure S4.** Pairwise phylogenetic distances and expression distances without echidna (R = 0.445, p = 1.09e-09).

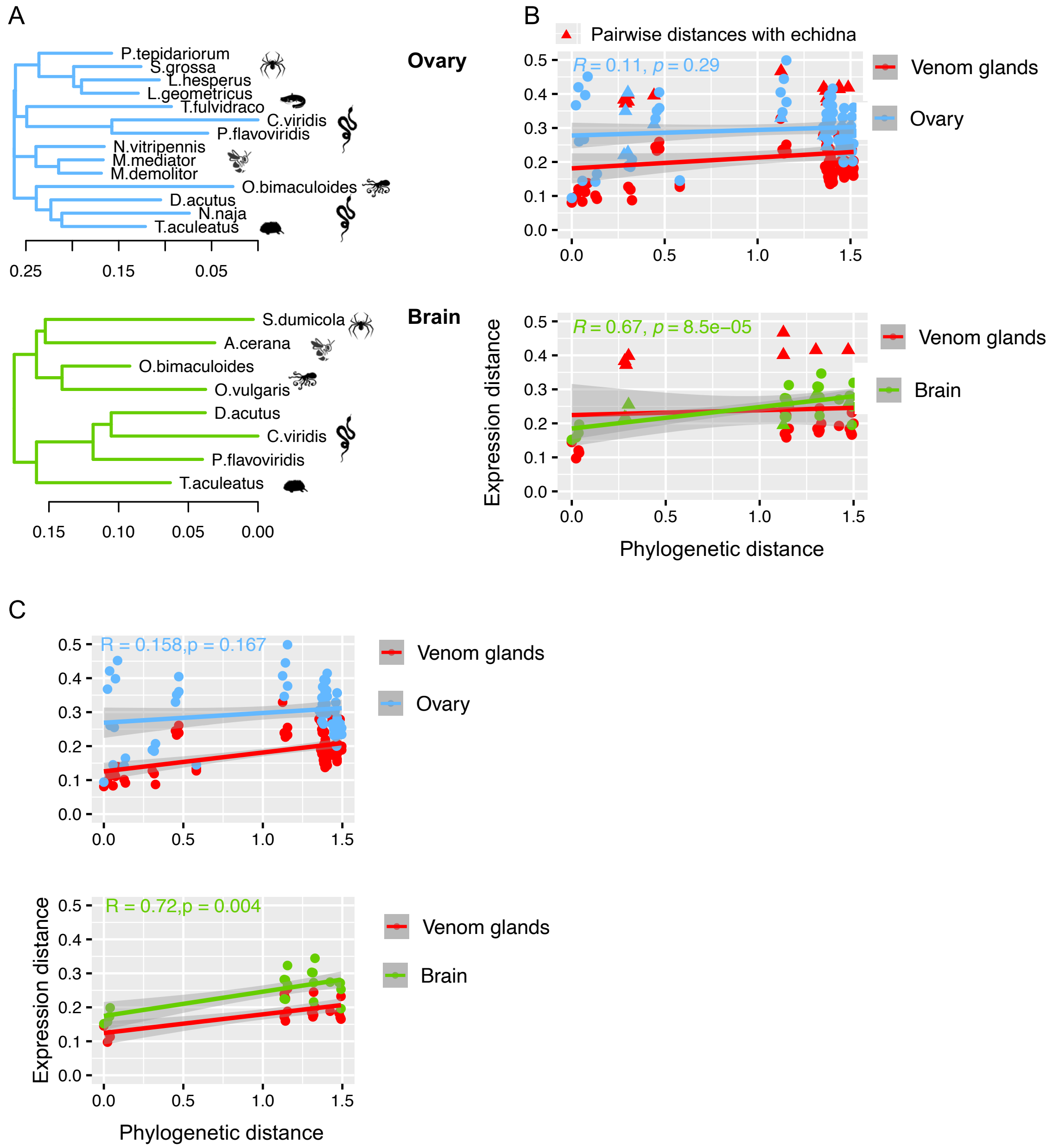


**Figure S5.** Divergence of tissue transcriptomes between species. A: Expression trees for the ovary and brain tissues. B: Sequence-based phylogenetic distances vs expression distances (1-Spearman coefficient) of ovary and brain. Pair distances between echidna and the other species are marked with triangles. C: As (B) but excluding pairwise comparisons with echidna.

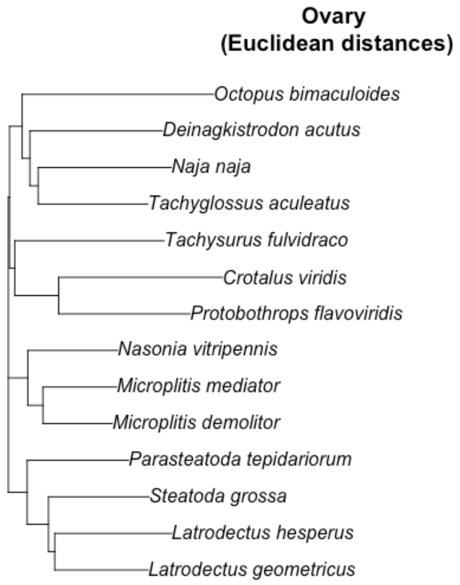


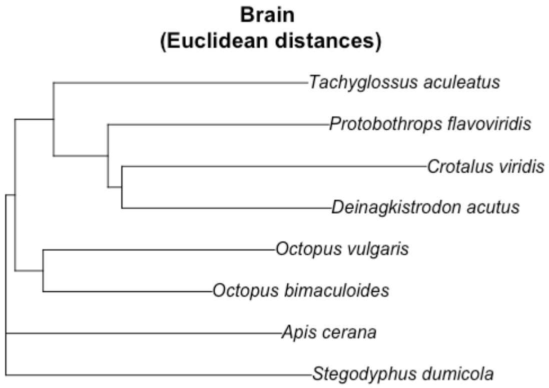


**Figure S6.** Ovary and brain expression trees based on Euclidean distance matrix.

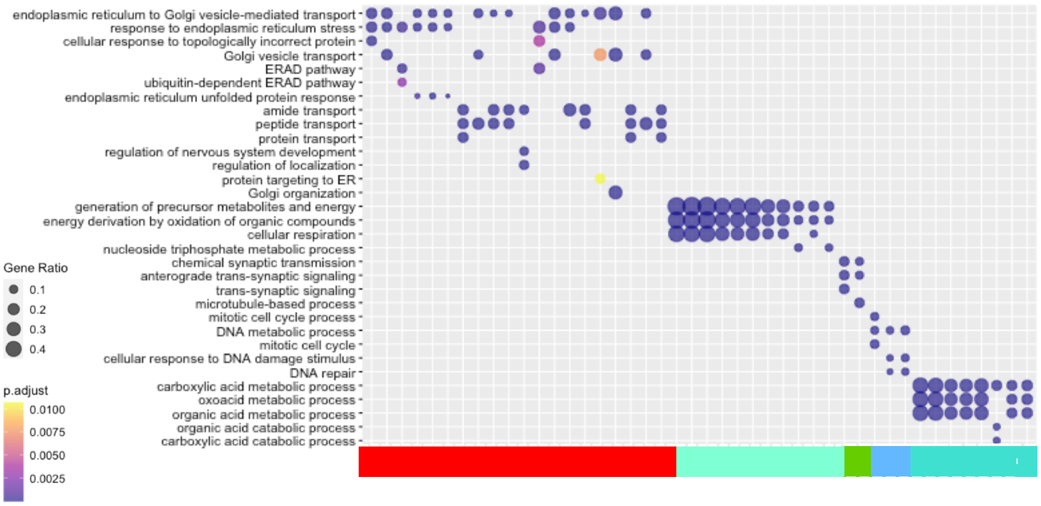


**Figure S7.** Enrichment of the top 3 biological process GO terms of the tissue-specific modules based on annotation of *Drosophila melanogaster* orthologs.

.

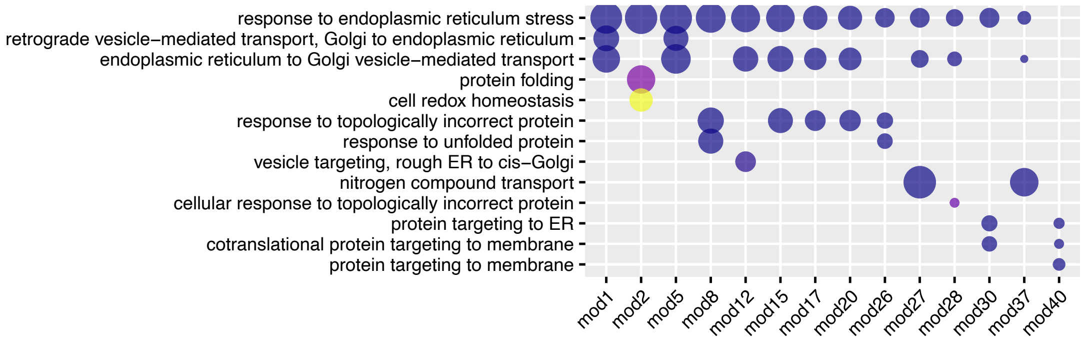


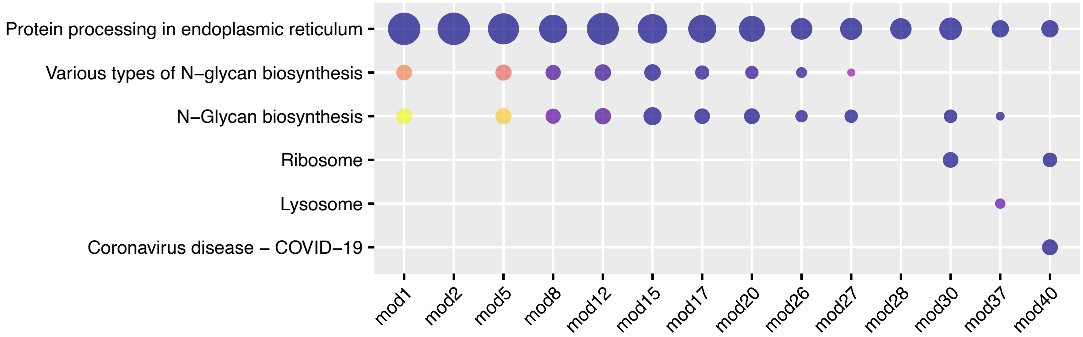


**Figure S8.** Enrichment of the top 3 biological process GO terms of the venom gland-specific modules obtained using the expression matrix based on transcripts selected randomly for each orthogroup.

**Table S1.** List of RNA-seq raw reads obtained from the European Nucleotide Archive (ENA).

| SRA study | Library type | SRA | Lineage | Species | Species code | Tissue | Tissue classification | Comments |
|---|---|---|---|---|---|---|---|---|
| SRP158111 | Paired | SRR7702281 | Fishes | *Tachysurus fulvidraco* | Tafu | Liver | Other organs | |
| SRP158111 | Paired | SRR7702282 | Fishes | *Tachysurus fulvidraco* | Tafu | Liver | Other organs | |
| SRP158111 | Paired | SRR7702283 | Fishes | *Tachysurus fulvidraco* | Tafu | Liver | Other organs | |
| SRP060326 | Paired | SRR2086959 | Fishes | *Tachysurus fulvidraco* | Tafu | Ovary | Ovary | |
| SRP057554 | Paired | SRR2002564 | Fishes | *Tachysurus fulvidraco* | Tafu | Venom gland | Venom glands | |
| SRP096985 | Paired | SRR5192547 | Flies | *Dasypogon diadema* | Dadi | Body tissue | Body tissues | |
| SRP096985 | Paired | SRR5192548 | Flies | *Dasypogon diadema* | Dadi | Body tissue | Body tissues | |
| SRP096985 | Paired | SRR7754487 | Flies | *Dasypogon diadema* | Dadi | Proboscis | Muscle tissues | |
| SRP096985 | Paired | SRR7754488 | Flies | *Dasypogon diadema* | Dadi | Proboscis | Muscle tissues | |
| SRP096985 | Paired | SRR7754485 | Flies | *Dasypogon diadema* | Dadi | Venom gland | Venom glands | |
| SRP096985 | Paired | SRR7754486 | Flies | *Dasypogon diadema* | Dadi | Venom gland | Venom glands | |
| SRP233233 | Single | SRR10530487 | Mammals | *Tachyglossus aculeatus* | Taac | Brain | Brain | |
| SRP233233 | Single | SRR10530493 | Mammals | *Tachyglossus aculeatus* | Taac | Brain | Brain | |
| SRP233233 | Single | SRR10530486 | Mammals | *Tachyglossus aculeatus* | Taac | Heart | Muscle tissues | |
| SRP233233 | Single | SRR10530492 | Mammals | *Tachyglossus aculeatus* | Taac | Heart | Muscle tissues | |
| SRP233233 | Single | SRR10530484 | Mammals | *Tachyglossus aculeatus* | Taac | Kidney | Other organs | |
| SRP233233 | Single | SRR10530488 | Mammals | *Tachyglossus aculeatus* | Taac | Kidney | Other organs | |
| SRP233233 | Single | SRR10530483 | Mammals | *Tachyglossus aculeatus* | Taac | Liver | Other organs | |
| SRP233233 | Single | SRR10530490 | Mammals | *Tachyglossus aculeatus* | Taac | Liver | Other organs | |
| SRP233233 | Single | SRR10530489 | Mammals | *Tachyglossus aculeatus* | Taac | Ovary | Ovary | |
| SRP027593 | Paired | SRR931704 | Mammals | *Tachyglossus aculeatus* | Taac | Venom gland | Venom glands | |
| SRP058882 | Paired | SRR2047122 | Octopi | *Octopus bimaculoides* | Ocbi | Viscera | Body tissues | hepatopancreas, kidney, heart |
| SRP058882 | Paired | SRR2048495 | Octopi | *Octopus bimaculoides* | Ocbi | Brain | Brain | subesophageal brain |
| SRP058882 | Paired | SRR2048496 | Octopi | *Octopus bimaculoides* | Ocbi | Brain | Brain | subesophageal brain |
| SRP058882 | Paired | SRR2048521 | Octopi | *Octopus bimaculoides* | Ocbi | Brain | Brain | supraesophageal brain |
| SRP058882 | Paired | SRR2048522 | Octopi | *Octopus bimaculoides* | Ocbi | Brain | Brain | supraesophageal brain |
| SRP058882 | Paired | SRR2045866 | Octopi | *Octopus bimaculoides* | Ocbi | Ovary | Ovary | |
| SRP058882 | Paired | SRR2047107 | Octopi | *Octopus bimaculoides* | Ocbi | Venom gland | Venom glands | posterior salivary gland |
| SRP154595 | Paired | SRR7548199 | Octopi | *Octopus vulgaris* | Ocvu | Brain | Brain | Hatchling, Vertical Lobe, Optic Lobe, Stellate Ganglion, Arm Cord, Superior Frontal Lobe |
| SRP144865 | Paired | SRR7130741 | Octopi | *Octopus vulgaris* | Ocvu | Venom gland | Venom glands | posterior salivary gland |
| SRP083064 | Paired | SRR6041834 | Scorpions | *Centruroides hentzi* | Cehe | Venom gland | Venom glands | mapped to *C. sculpturatus* |

| | | | | | | | | |
|---|---|---|---|---|---|---|---|---|
| SRP083064 | Paired | SRR6041835 | Scorpions | *Centruroides hentzi* | Cehe | Venom gland | Venom glands | mapped to *C. sculpturatus* |
| SRP064632 | Paired | SRR3061371 | Scorpions | *Mesobuthus martensii* | Mema | Abdomen | Body tissues | |
| SRP064632 | Paired | SRR3984610 | Scorpions | *Mesobuthus martensii* | Mema | Abdomen | Body tissues | |
| SRP064632 | Paired | SRR2592319 | Scorpions | *Mesobuthus martensii* | Mema | Cephalothorax | Body tissues | |
| SRP064632 | Paired | SRR3056832 | Scorpions | *Mesobuthus martensii* | Mema | Cephalothorax | Body tissues | |
| SRP064632 | Paired | SRR3984597 | Scorpions | *Mesobuthus martensii* | Mema | Cephalothorax | Body tissues | |
| SRP064632 | Paired | SRR2592957 | Scorpions | *Mesobuthus martensii* | Mema | Muscle | Muscle tissues | |
| SRP064632 | Paired | SRR3061373 | Scorpions | *Mesobuthus martensii* | Mema | Muscle | Muscle tissues | |
| SRP064632 | Paired | SRR3984642 | Scorpions | *Mesobuthus martensii* | Mema | Muscle | Muscle tissues | |
| SRP064632 | Paired | SRR2592960 | Scorpions | *Mesobuthus martensii* | Mema | Venom gland | Venom glands | |
| SRP064632 | Paired | SRR3061379 | Scorpions | *Mesobuthus martensii* | Mema | Venom gland | Venom glands | |
| SRP064632 | Paired | SRR3984663 | Scorpions | *Mesobuthus martensii* | Mema | Venom gland | Venom glands | |
| SRP065240 | Paired | SRR2790072 | Scorpions | *Mesobuthus martensii* | Mema | Venom gland | Venom glands | |
| SRP065240 | Paired | SRR2791632 | Scorpions | *Mesobuthus martensii* | Mema | Venom gland | Venom glands | after electric shock |
| SRP087425 | Paired | SRR4188636 | Scorpions | *Mesobuthus martensii* | Mema | Venom gland | Venom glands | |
| SRP150951 | Paired | SRR7401995 | Snakes | *Crotalus viridis* | Crvi | Brain | Brain | |
| SRP150951 | Paired | SRR7401993 | Snakes | *Crotalus viridis* | Crvi | Muscle | Muscle tissues | |
| SRP150951 | Paired | SRR7401990 | Snakes | *Crotalus viridis* | Crvi | Accessory venom gland | Other glands | |
| SRP150951 | Paired | SRR7401996 | Snakes | *Crotalus viridis* | Crvi | Rictal gland | Other glands | |
| SRP150951 | Paired | SRR7401980 | Snakes | *Crotalus viridis* | Crvi | Kidney | Other organs | |
| SRP150951 | Paired | SRR7401984 | Snakes | *Crotalus viridis* | Crvi | Kidney | Other organs | |
| SRP150951 | Paired | SRR7401985 | Snakes | *Crotalus viridis* | Crvi | Kidney | Other organs | |
| SRP150951 | Paired | SRR7401978 | Snakes | *Crotalus viridis* | Crvi | Liver | Other organs | |
| SRP150951 | Paired | SRR7401982 | Snakes | *Crotalus viridis* | Crvi | Liver | Other organs | |
| SRP150951 | Paired | SRR7401983 | Snakes | *Crotalus viridis* | Crvi | Liver | Other organs | |
| SRP150951 | Paired | SRR7401986 | Snakes | *Crotalus viridis* | Crvi | Pancreas | Other organs | |
| SRP150951 | Paired | SRR7402009 | Snakes | *Crotalus viridis* | Crvi | Pancreas | Other organs | |
| SRP150951 | Paired | SRR7401997 | Snakes | *Crotalus viridis* | Crvi | Ovary | Ovary | |
| SRP150951 | Paired | SRR7401989 | Snakes | *Crotalus viridis* | Crvi | Venom gland | Venom glands | unextracted |
| SRP150951 | Paired | SRR7402004 | Snakes | *Crotalus viridis* | Crvi | Venom gland | Venom glands | 1day post milking |
| SRP150951 | Paired | SRR7402005 | Snakes | *Crotalus viridis* | Crvi | Venom gland | Venom glands | 3days post milkng |
| SRP071324 | Paired | SRR3212849 | Snakes | *Deinagkistrodon acutus* | Deac | Brain | Brain | |
| SRP071324 | Paired | SRR3212852 | Snakes | *Deinagkistrodon acutus* | Deac | Brain | Brain | |
| SRP071324 | Paired | SRR3212850 | Snakes | *Deinagkistrodon acutus* | Deac | Liver | Other organs | |
| SRP071324 | Paired | SRR3212854 | Snakes | *Deinagkistrodon acutus* | Deac | Liver | Other organs | |
| SRP071324 | Paired | SRR3212847 | Snakes | *Deinagkistrodon acutus* | Deac | Ovary | Ovary | |

| | | | | | | | | |
|---|---|---|---|---|---|---|---|---|
| SRP071324 | Paired | SRR3212848 | Snakes | *Deinagkistrodon acutus* | Deac | Venom gland | Venom glands | |
| SRP071324 | Paired | SRR3212851 | Snakes | *Deinagkistrodon acutus* | Deac | Venom gland | Venom glands | |
| SRP188892 | Paired | SRR8754985 | Snakes | *Naja naja* | Nana | Heart | Muscle tissues | |
| SRP188892 | Single | SRR8754972 | Snakes | *Naja naja* | Nana | Salivary gland | Other glands | |
| SRP188892 | Single | SRR8754966 | Snakes | *Naja naja* | Nana | Kidney | Other organs | |
| SRP188892 | Single | SRR8754970 | Snakes | *Naja naja* | Nana | Kidney | Other organs | |
| SRP188892 | Paired | SRR8754989 | Snakes | *Naja naja* | Nana | Liver | Other organs | |
| SRP188892 | Single | SRR8754967 | Snakes | *Naja naja* | Nana | Liver | Other organs | |
| SRP188892 | Paired | SRR8754987 | Snakes | *Naja naja* | Nana | Pancreas | Other organs | |
| SRP188892 | Single | SRR8754974 | Snakes | *Naja naja* | Nana | Pancreas | Other organs | |
| SRP188892 | Single | SRR8754975 | Snakes | *Naja naja* | Nana | Pancreas | Other organs | |
| SRP188892 | Single | SRR8754969 | Snakes | *Naja naja* | Nana | Ovary | Ovary | |
| SRP188892 | Paired | SRR8754977 | Snakes | *Naja naja* | Nana | Venom gland | Venom glands | |
| SRP188892 | Paired | SRR8754986 | Snakes | *Naja naja* | Nana | Venom gland | Venom glands | |
| SRP188892 | Single | SRR8754980 | Snakes | *Naja naja* | Nana | Venom gland | Venom glands | |
| SRP188892 | Single | SRR8754981 | Snakes | *Naja naja* | Nana | Venom gland | Venom glands | |
| DRP004366 | Paired | DRR125545 | Snakes | *Protobothrops flavoviridis* | Prfl | Brain | Brain | |
| DRP004366 | Paired | DRR125549 | Snakes | *Protobothrops flavoviridis* | Prfl | Brain | Brain | |
| DRP004366 | Paired | DRR125558 | Snakes | *Protobothrops flavoviridis* | Prfl | Heart | Muscle tissues | |
| DRP004366 | Paired | DRR125560 | Snakes | *Protobothrops flavoviridis* | Prfl | Muscle | Muscle tissues | |
| DRP004366 | Paired | DRR125553 | Snakes | *Protobothrops flavoviridis* | Prfl | Kidney | Other organs | |
| DRP004366 | Paired | DRR125552 | Snakes | *Protobothrops flavoviridis* | Prfl | Liver | Other organs | |
| DRP004366 | Paired | DRR125554 | Snakes | *Protobothrops flavoviridis* | Prfl | Pancreas | Other organs | |
| DRP004366 | Paired | DRR125559 | Snakes | *Protobothrops flavoviridis* | Prfl | Ovary | Ovary | |
| DRP004366 | Paired | DRR024168 | Snakes | *Protobothrops flavoviridis* | Prfl | Venom gland | Venom glands | |
| DRP004366 | Paired | DRR125542 | Snakes | *Protobothrops flavoviridis* | Prfl | Venom gland | Venom glands | |
| DRP004366 | Paired | DRR125548 | Snakes | *Protobothrops flavoviridis* | Prfl | Venom gland | Venom glands | |
| SRP100704 | Paired | SRR5285084 | Spiders | *Latrodectus geometricus* | Lage | Cephalothorax | Body tissues | |
| SRP100704 | Paired | SRR5285085 | Spiders | *Latrodectus geometricus* | Lage | Cephalothorax | Body tissues | |
| SRP100704 | Paired | SRR5285088 | Spiders | *Latrodectus geometricus* | Lage | Silk gland | Other glands | original tissue name: MAJAMPSG |
| SRP100704 | Paired | SRR5285093 | Spiders | *Latrodectus geometricus* | Lage | Silk gland | Other glands | original tissue name: MINAMPSG |
| SRP100704 | Paired | SRR5285097 | Spiders | *Latrodectus geometricus* | Lage | Silk gland | Other glands | original tissue name: TUBULSG |
| SRP100704 | Paired | SRR5285094 | Spiders | *Latrodectus geometricus* | Lage | Ovary | Ovary | |
| SRP100704 | Paired | SRR5285095 | Spiders | *Latrodectus geometricus* | Lage | Ovary | Ovary | |
| SRP100704 | Paired | SRR5285099 | Spiders | *Latrodectus geometricus* | Lage | Venom gland | Venom glands | |
| SRP100704 | Paired | SRR5285100 | Spiders | *Latrodectus geometricus* | Lage | Venom gland | Venom glands | |
| SRP040992 | Paired | SRR1219650 | Spiders | *Latrodectus hesperus* | Lahe | Cephalothorax | Body tissues | |

| | | | | | | | | |
|---|---|---|---|---|---|---|---|---|
| SRP040992 | Paired | SRR1219651 | Spiders | *Latrodectus hesperus* | Lahe | Cephalothorax | Body tissues | |
| SRP100704 | Paired | SRR5285104 | Spiders | *Latrodectus hesperus* | Lahe | Cephalothorax | Body tissues | |
| SRP100704 | Paired | SRR5285109 | Spiders | *Latrodectus hesperus* | Lahe | Silk gland | Other glands | original tissue name: MAJAMPSG |
| SRP100704 | Paired | SRR5285113 | Spiders | *Latrodectus hesperus* | Lahe | Silk gland | Other glands | original tissue name: MINAMPSG |
| SRP100704 | Paired | SRR5285119 | Spiders | *Latrodectus hesperus* | Lahe | Silk gland | Other glands | original tissue name: TUBULSG |
| SRP100704 | Paired | SRR5285114 | Spiders | *Latrodectus hesperus* | Lahe | Ovary | Ovary | |
| SRP100704 | Paired | SRR5285115 | Spiders | *Latrodectus hesperus* | Lahe | Ovary | Ovary | |
| SRP040992 | Paired | SRR1219652 | Spiders | *Latrodectus hesperus* | Lahe | Venom gland | Venom glands | |
| SRP100704 | Paired | SRR5285121 | Spiders | *Latrodectus hesperus* | Lahe | Venom gland | Venom glands | |
| SRP100704 | Paired | SRR5285122 | Spiders | *Latrodectus hesperus* | Lahe | Venom gland | Venom glands | |
| SRP100704 | Paired | SRR5285123 | Spiders | *Latrodectus hesperus* | Lahe | Venom gland | Venom glands | |
| SRP188917 | Paired | SRR8755629 | Spiders | *Parasteatoda tepidariorum* | Pate | Cephalothorax | Body tissues | |
| SRP188917 | Paired | SRR8755630 | Spiders | *Parasteatoda tepidariorum* | Pate | Cephalothorax | Body tissues | |
| SRP188917 | Paired | SRR8755627 | Spiders | *Parasteatoda tepidariorum* | Pate | Silk gland | Other glands | |
| SRP188917 | Paired | SRR8755628 | Spiders | *Parasteatoda tepidariorum* | Pate | Silk gland | Other glands | |
| SRP055769 | Paired | SRR1824489 | Spiders | *Parasteatoda tepidariorum* | Pate | Ovary | Ovary | |
| SRP188917 | Paired | SRR8755633 | Spiders | *Parasteatoda tepidariorum* | Pate | Ovary | Ovary | |
| SRP188917 | Paired | SRR8755634 | Spiders | *Parasteatoda tepidariorum* | Pate | Ovary | Ovary | |
| SRP095650 | Paired | SRR5131058 | Spiders | *Parasteatoda tepidariorum* | Pate | Venom gland | Venom glands | |
| SRP188917 | Paired | SRR8755631 | Spiders | *Parasteatoda tepidariorum* | Pate | Venom gland | Venom glands | |
| SRP188917 | Paired | SRR8755632 | Spiders | *Parasteatoda tepidariorum* | Pate | Venom gland | Venom glands | |
| SRP100704 | Paired | SRR5285126 | Spiders | *Steatoda grossa* | Stgr | Cephalothorax | Body tissues | |
| SRP100704 | Paired | SRR5285127 | Spiders | *Steatoda grossa* | Stgr | Cephalothorax | Body tissues | |
| SRP100704 | Paired | SRR5285130 | Spiders | *Steatoda grossa* | Stgr | Silk gland | Other glands | original tissue name: MAJAMPSG |
| SRP100704 | Paired | SRR5285134 | Spiders | *Steatoda grossa* | Stgr | Silk gland | Other glands | original tissue name: MINAMPSG |
| SRP100704 | Paired | SRR5285139 | Spiders | *Steatoda grossa* | Stgr | Silk gland | Other glands | original tissue name: TUBULSG |
| SRP100704 | Paired | SRR5285135 | Spiders | *Steatoda grossa* | Stgr | Ovary | Ovary | |
| SRP100704 | Paired | SRR5285136 | Spiders | *Steatoda grossa* | Stgr | Ovary | Ovary | |
| SRP100704 | Paired | SRR5285141 | Spiders | *Steatoda grossa* | Stgr | Venom gland | Venom glands | |
| SRP100704 | Paired | SRR5285142 | Spiders | *Steatoda grossa* | Stgr | Venom gland | Venom glands | |
| SRP223964 | Paired | SRR10216521 | Spiders | *Stegodyphus dumicola* | Stdu | Abdomen | Body tissues | |
| SRP223964 | Paired | SRR10216524 | Spiders | *Stegodyphus dumicola* | Stdu | Abdomen | Body tissues | |
| SRP223964 | Paired | SRR10216525 | Spiders | *Stegodyphus dumicola* | Stdu | Abdomen | Body tissues | |
| SRP223964 | Paired | SRR10216518 | Spiders | *Stegodyphus dumicola* | Stdu | Brain | Brain | |
| SRP223964 | Paired | SRR10216519 | Spiders | *Stegodyphus dumicola* | Stdu | Brain | Brain | |
| SRP223964 | Paired | SRR10216520 | Spiders | *Stegodyphus dumicola* | Stdu | Brain | Brain | |
| SRP223964 | Paired | SRR10216515 | Spiders | *Stegodyphus dumicola* | Stdu | Leg | Muscle tissues | |

| | | | | | | | | |
|---|---|---|---|---|---|---|---|---|
| SRP223964 | Paired | SRR10216516 | Spiders | *Stegodyphus dumicola* | Stdu | Leg | Muscle tissues | |
| SRP223964 | Paired | SRR10216517 | Spiders | *Stegodyphus dumicola* | Stdu | Leg | Muscle tissues | |
| SRP223964 | Paired | SRR10216514 | Spiders | *Stegodyphus dumicola* | Stdu | Venom gland | Venom glands | |
| SRP223964 | Paired | SRR10216522 | Spiders | *Stegodyphus dumicola* | Stdu | Venom gland | Venom glands | |
| SRP223964 | Paired | SRR10216523 | Spiders | *Stegodyphus dumicola* | Stdu | Venom gland | Venom glands | |
| SRP043101 | Paired | SRR1380970 | Wasps | *Apis cerana* | Apce | Brain | Brain | |
| SRP043101 | Paired | SRR1380979 | Wasps | *Apis cerana* | Apce | Hypopharingeal gland | Other glands | |
| SRP043101 | Paired | SRR1406762 | Wasps | *Apis cerana* | Apce | Venom gland | Venom glands | |
| SRP028964 | Paired | SRR955015 | Wasps | *Microplitis demolitor* | Mide | Ovary | Ovary | |
| SRP028964 | Paired | SRR955397 | Wasps | *Microplitis demolitor* | Mide | Venom gland | Venom glands | |
| SRP096668 | Paired | SRR5177962 | Wasps | *Microplitis mediator* | Mime | Ovary | Ovary | mapped to *M. demolitor* |
| SRP114339 | Paired | SRR5885434 | Wasps | *Microplitis mediator* | Mime | Ovary | Ovary | mapped to *M. demolitor* |
| SRP114339 | Paired | SRR5885444 | Wasps | *Microplitis mediator* | Mime | Ovary | Ovary | mapped to *M. demolitor* |
| SRP114339 | Paired | SRR5885445 | Wasps | *Microplitis mediator* | Mime | Ovary | Ovary | mapped to *M. demolitor* |
| SRP096668 | Paired | SRR5177969 | Wasps | *Microplitis mediator* | Mime | Venom gland | Venom glands | mapped to *M. demolitor* |
| SRP114339 | Paired | SRR5885438 | Wasps | *Microplitis mediator* | Mime | Venom gland | Venom glands | mapped to *M. demolitor* |
| SRP114339 | Paired | SRR5885439 | Wasps | *Microplitis mediator* | Mime | Venom gland | Venom glands | mapped to *M. demolitor* |
| SRP114339 | Paired | SRR5885440 | Wasps | *Microplitis mediator* | Mime | Venom gland | Venom glands | mapped to *M. demolitor* |
| SRP067692 | Paired | SRR3046456 | Wasps | *Nasonia vitripennis* | Navi | Ovary | Ovary | |
| SRP067692 | Paired | SRR3046457 | Wasps | *Nasonia vitripennis* | Navi | Ovary | Ovary | |
| SRP067692 | Paired | SRR3046458 | Wasps | *Nasonia vitripennis* | Navi | Ovary | Ovary | |
| SRP067692 | Paired | SRR3046453 | Wasps | *Nasonia vitripennis* | Navi | Venom gland | Venom glands | |
| SRP067692 | Paired | SRR3046454 | Wasps | *Nasonia vitripennis* | Navi | Venom gland | Venom glands | |
| SRP067692 | Paired | SRR3046455 | Wasps | *Nasonia vitripennis* | Navi | Venom gland | Venom glands | |

**Table S2.** List of species used for mapping in OrthoDB at the Metazoa node.

| Lineage | Species | OrthoDB reference species |
|---|---|---|
| **Wasps** | *Nasonia vitripennis, Microplitis demolitor, Microplitis mediator* | Microplitis demolitor, genome GCF_000572035.2 |
| | | Nasonia vitripennis, genome GCF_000002325.3 |
| | *Apis cerana* | Apis mellifera, genome GCF_000002195.4 |
| | | Polistes canadensis, genome GCF_001313835.1 |
| | | Nasonia vitripennis, genome GCF_000002325.3 |
| **Flies** | *Dasypogon diadema* | Mayetiola destructor, genome 0 |
| | | Drosophila grimshawi, genome 0 |
| | | Drosophila melanogaster, genome dmel_r6.19 |
| | | Drosophila obscura, genome GCF_002217835.1 |
| | | Lucilia cuprina, genome GCF_000699065.1 |
| | | Anopheles darlingi, genome 0 |
| | | Anopheles gambiae, genome 0 |
| | | Aedes aegypti, genome 0 |
| | | Culex quinquefasciatus, genome 0 |
| **Spiders and Scorpions** | *Parasteatoda tepidariorum, Stegodyphus dumicola, Steatoda grossa, Latrodectus geometricus, Latrodectus hesperus, Centruroides hentzi, Mesobuthus martensii* | Parasteatoda tepidariorum, genome GCF_000365465.2 |
| | | Stegodyphus mimosarum, genome GCA_000611955.2 |
| | | Tetranychus urticae, genome GCF_000239435.1 |
| | | Tropilaelaps mercedesae, genome GCA_002081605.1 |
| | | Varroa jacobsoni, genome GCF_002532875.1 |
| | | Euroglyphus maynei, genome GCA_002135145.1 |
| | | Galendromus occidentalis, genome GCF_000255335.1 |
| | | Ixodes scapularis, genome 0 |
| | | Sarcoptes scabiei, genome 0 |
| | | Centruroides sculpturatus, genome GCF_000671375.1 |
| **Octopi** | *Octopus bimaculoides, Octopus vulgaris* | Octopus bimaculoides, genome GCF_001194135.1 |
| **Fishes** | *Tachysurus fulvidraco* | Astyanax mexicanus, genome GCF_000372685.2 |
| | | Danio rerio, genome GCF_000002035.6 |
| | | Ictalurus punctatus, genome GCF_001660625.1 |
| | | Pygocentrus nattereri, genome GCF_001682695.1 |
| **Mammals** | *Tachyglossus aculeatus* | Ornithorhynchus anatinus, genome GCF_000002275.2 |
| **Snakes** | *Naja naja Deinagkistrodon acutus, Protobothrops flavoviridis, Crotalus viridis* | Anolis carolinensis, genome GCF_000090745.1 |
| | | Gekko japonicus, genome GCF_001447785.1 |
| | | Protobothrops mucrosquamatus, genome GCF_001527695.x |
| | | Python bivittatus, genome GCF_000186305.1 |
| | | Thamnophis sirtalis, genome GCF_001077635.1 |

**Legends for Datasets S1 to S9**

**Dataset S1 (separate file).** Expression matrix of $log_2$-transformed, quantile normalized Transcripts Per Million (TPM) values of the 2,528 orthologs shared among all species. Species abbreviations are listed in Table S1.

**Dataset S2 (separate file).** Orthogroup weights of gene modules identified with isa clustering.

**Dataset S3 (separate file).** Sample weights of gene modules identified with isa clustering. Species abbreviations are listed in Table S1.

**Dataset S4 (separate file).** Results of biological process GO terms enrichment analysis of gene modules based on human annotation.

**Dataset S5 (separate file).** Results of KEGG pathway enrichment analysis of gene modules based on human annotation.

**Dataset S6 (separate file).** Results of KEGG pathway enrichment analyses of significantly upregulated genes. Each species was analyzed separately. The species used for the annotation are reported ('Reference_species').

**Dataset S7 (separate file).** Results of GO terms enrichment analyses of significantly upregulated genes. Each species was analyzed separately.

**Dataset S8 (separate file).** List of transcription factors expressed in venom glands. For each species is reported the logarithmic fold-change (logFC), and the logarithmic Count per Million (logCPM). Species abbreviations are listed in Table S1.

**Dataset S9 (separate file).** List of representative sequences for each orthogroup in each species selected as the transcript with the highest expression in venom gland samples.
